# Experimental Evidence for HIV-Associated Phenotypic Reprogramming of CD4□ T Cells: An Exploratory Immunological Study

**DOI:** 10.64898/2026.07.18.739371

**Authors:** Sherif Salah, Khalid F. Kassem, Mohamed Sherif, Ashraf M. Abu-Seida

## Abstract

**Background:** Human immunodeficiency virus type 1 (HIV-1) infection is characterized by progressive immune dysfunction, classically attributed to depletion of CD4□ T lymphocytes. Despite decades of research, the mechanisms underlying the functional deterioration of cellular immunity remain incompletely understood. We hypothesized that, in addition to quantitative CD4□ T-cell loss, HIV infection may induce phenotypic reprogramming of CD4□ T cells, resulting in altered surface-marker expression and impaired immunological function.

**Methods:** Peripheral blood samples were obtained from untreated HIV-positive individuals and healthy controls. Flow cytometric immunophenotyping, recombinant HIV-1 p17 stimulation assays, immunofluorescence imaging, ELISPOT analysis of interferon-γ secretion, and CD4□/CD8□conjugation assays were performed to investigate dynamic changes in T-cell phenotype and function following viral protein exposure.

**Results:** HIV-positive samples demonstrated progressive reductions in CD4□ T-cell counts accompanied by corresponding increases in CD8□ T-cell populations following p17 stimulation, while the combined CD4□/CD8□ T-cell count remained relatively stable. Immunofluorescence analyses identified cells exhibiting simultaneous CD4- and CD8-associated marker expression after prolonged incubation. Functional analyses further demonstrated markedly reduced interferon-γ secretion in the newly identified cell population compared with conventional CD8□ T cells. Conjugation assays additionally suggested altered cellular interaction patterns between reprogrammed cells and target CD4□lymphocytes.

**Conclusions:** These findings support the hypothesis that chronic HIV infection may be associated with phenotypic reprogramming of CD4□ T cells rather than simple quantitative depletion alone. Although the underlying molecular mechanisms remain to be established, this exploratory model provides an alternative framework for investigating HIV-associated immune dysfunction and may stimulate future studies using lineage-tracing, single-cell transcriptomics, and epigenetic profiling to evaluate this proposed mechanism.

## 1. Introduction

Human immunodeficiency virus type 1 (HIV-1) remains one of the most important chronic viral infections worldwide, affecting millions of individuals despite remarkable advances in antiretroviral therapy (ART). Although ART effectively suppresses viral replication and has substantially improved patient survival, it does not eliminate persistent viral reservoirs or completely restore normal immune function [1–4]. Many treated individuals continue to exhibit immune activation, chronic inflammation, and immunological abnormalities despite sustained virological suppression, indicating that mechanisms beyond active viral replication contribute to HIV pathogenesis [2–4]. Understanding these mechanisms remains essential for developing therapeutic strategies capable of achieving durable immune restoration and, ultimately, a functional cure.

The progressive decline in circulating CD4□ T lymphocytes has long been considered the immunological hallmark of HIV infection. Current models attribute this decline to multiple complementary mechanisms, including direct viral cytopathic effects, activation-induced apoptosis, pyroptosis of abortively infected bystander cells, chronic immune activation, microbial translocation resulting from gastrointestinal barrier disruption, and progressive immune exhaustion [5–11]. Together, these processes explain much of the quantitative depletion of helper T cells observed during chronic infection. Nevertheless, several aspects of HIV-associated immune dysfunction remain incompletely understood. In particular, the relationship among progressive CD4□ T-cell loss, expansion, and persistent activation of CD8□ T-cell populations, and deterioration of immune competence may not be fully represented by absolute circulating lymphocyte counts alone [12–18].

CD4□ and CD8□ T lymphocytes function as highly coordinated components of adaptive immunity. CD4□ T cells orchestrate antiviral responses through cytokine production, activation of antigen-presenting cells, regulation of B-cell maturation, and provision of essential support for effective cytotoxic and memory T-cell responses. In contrast, CD8□ T cells recognize and eliminate infected host cells through perforin- and granzyme-mediated cytotoxicity while producing antiviral cytokines that contribute to viral control [12–18]. The biological activity of each population therefore depends not only on its independent effector functions but also on continuous reciprocal communication between helper and cytotoxic lymphocytes. Consequently, disruption of either the quantitative distribution or qualitative functional state of these populations may profoundly affect immune homeostasis.

Recent advances in high-dimensional immunophenotyping and single-cell analysis have shown that apparently uniform CD4□ and CD8□ T-cell populations contain diverse transcriptional and functional states, including exhausted, interferon-responsive, metabolically altered, and transitional cellular populations [19–23]. Mature lymphocytes are therefore increasingly recognized as dynamic populations capable of adapting their transcriptional programs, metabolic activity, receptor expression, and functional properties in response to persistent environmental signals.

This adaptive capacity is commonly described as immune-cell plasticity or phenotypic reprogramming. Experimental studies have shown that mature CD4□ T cells can modify lineage-associated transcriptional networks, cytokine programs, epigenetic states, and effector functions under defined environmental conditions [24–31]. These changes can involve reciprocal regulation of lineage-associated factors such as ThPOK and Runx3, acquisition of cytotoxic properties, and remodeling of chromatin accessibility without necessarily implying irreversible genomic mutation or complete conversion into a different lymphocyte lineage [24–31]. Phenotypic reprogramming therefore provides a biologically plausible framework through which persistent inflammatory or antigenic stimulation could modify cellular identity and function while preserving lineage continuity.

Within the context of HIV infection, extensive investigation has focused on immune activation, exhaustion, transcriptional dysregulation, metabolic disturbance, and the heterogeneous cellular composition of persistent viral reservoirs [2–4,14–23,41–44]. However, comparatively less attention has been directed toward whether prolonged HIV-associated stimulation may induce broader phenotypic remodeling of CD4□ T cells beyond progressive numerical depletion or classical exhaustion. Such remodeling could potentially contribute to immune dysfunction by altering receptor expression and biological behavior without requiring immediate cellular elimination. Exploring this possibility does not challenge established mechanisms of HIV pathogenesis; rather, it considers whether qualitative phenotypic alterations may coexist with the well-recognized quantitative loss and functional impairment of helper T cells.

Extracellular HIV-1 p17 matrix protein has been reported to retain biological activity and to influence lymphocyte activation, inflammatory cytokine production, dendritic-cell function, chemokine-receptor signaling, and monocyte/macrophage responses [32–38]. Accordingly, recombinant p17 was selected as a controlled ex vivo stimulus for investigating whether exposure to an HIV-associated structural protein produces measurable changes in T-cell immunophenotype and function. Importantly, p17 stimulation is not equivalent to infection with replication-competent HIV-1; it represents an experimental model for examining one component of HIV-associated cellular signaling.

The present exploratory study therefore investigated an alternative immunological model in which HIV-associated stimulation may promote phenotypic reprogramming of CD4□ T cells, resulting in altered surface-marker expression accompanied by functional changes that could contribute to immune dysregulation. To explore this hypothesis, we combined flow-cytometric immunophenotyping, recombinant HIV-1 p17 stimulation, immunofluorescence microscopy, interferon-γ enzyme-linked immunospot analysis, and cellular conjugation experiments using samples obtained from untreated HIV-positive individuals and healthy controls (Supplementary S1). Rather than proposing a replacement for current models of HIV immunopathogenesis, this study sought to determine whether the observed experimental findings support an additional, hypothesis-generating biological framework requiring validation by contemporary molecular and single-cell approaches.

### 1.1. Scientific Rationale, Knowledge Gap, and Study Hypothesis

Despite substantial progress in defining the molecular and cellular mechanisms underlying HIV infection, the relationship between lymphocyte abundance, cellular identity, and immune competence remains incompletely understood. Some individuals experience incomplete immune recovery despite sustained virological suppression, whereas clinical and functional outcomes may differ among individuals with similar circulating CD4□ T-cell counts [1–4,19–23]. These observations indicate that absolute lymphocyte numbers alone may not capture the full biological complexity of HIV-associated immune dysfunction.

Systems-level and single-cell studies have shown that immune-cell populations that appear numerically preserved may nevertheless display major alterations in transcriptional activity, metabolic regulation, exhaustion state, cytokine responsiveness, and effector function [19– 23,41–44]. Persistent antigenic exposure, inflammatory stimulation, and prolonged cytokine signaling can also induce epigenetic and transcriptional remodeling of differentiated T-cell populations [24–31]. Whether a related process contributes to the altered CD4□/CD8□immunophenotypic patterns observed following HIV-associated stimulation remains uncertain.

The present study was motivated by preliminary observations that could not be interpreted solely through immediate cellular loss. Following ex vivo exposure of samples from untreated HIV-positive participants to recombinant HIV-1 p17, progressive changes were observed in detectable CD4□ and CD8□ T-cell populations. These quantitative changes were accompanied by altered immunofluorescence staining patterns and functional readouts. Because extracellular p17 can exert immunomodulatory activity independently of complete viral replication [32–38], these observations prompted investigation of whether p17-associated signaling could induce an altered T-cell phenotypic state.

We therefore hypothesized that HIV-associated stimulation may induce phenotypic reprogramming in a subset of CD4□ T lymphocytes, producing progressive modification of their immunophenotypic profile together with impairment of normal helper-cell functions. In this manuscript, **phenotypic reprogramming** refers to experimentally observed alterations in surface-marker expression and biological behavior. It does not imply proven genetic mutation, irreversible lineage conversion, or definitive conversion of a CD4□ T cell into a conventional CD8□ T cell. Studies of mature T-cell plasticity nevertheless provide a conceptual basis for investigating such altered cellular states [24–31].

To evaluate this hypothesis, we combined longitudinal flow-cytometric immunophenotyping, recombinant HIV-1 p17 stimulation, dual-marker immunofluorescence microscopy, interferon-γ enzyme-linked immunospot analysis, and cellular interaction assays. The primary objective was to determine whether the convergence of these experimental findings supports the presence of an altered CD4-associated immunophenotypic and functional state following p17 stimulation. The proposed model is hypothesis-generating and requires validation through high-parameter flow cytometry, lineage-associated transcription-factor analysis, cell sorting, single-cell RNA sequencing, T-cell receptor clonotyping, and epigenomic profiling [19–31].

## 2. Materials and Methods

### 2.1 Study Design

This study was designed as an exploratory immunological investigation to evaluate whether chronic HIV-1 infection is associated with phenotypic reprogramming of CD4□ T lymphocytes and to examine the potential functional consequences of this proposed process. The experimental strategy combined quantitative immunophenotyping with complementary functional assays to investigate dynamic changes in T-cell phenotype following controlled ex vivo stimulation with recombinant HIV-1 proteins.

The study consisted of two principal components. The first evaluated longitudinal changes in CD4□ and CD8□ T-cell populations using flow cytometric immunophenotyping before and after stimulation with recombinant HIV-1 p17 matrix protein. The second employed complementary laboratory techniques, including immunofluorescence microscopy, interferon-γ enzyme-linked immunospot (ELISPOT) analysis, and cellular conjugation assays, to investigate alterations in surface-marker expression and functional behavior that accompanied the observed immunophenotypic changes.

Rather than testing an established biological mechanism, the study was designed to generate experimental evidence supporting or refuting an alternative hypothesis of HIV-associated immune dysregulation based on phenotypic reprogramming of CD4□ T lymphocytes. Accordingly, all findings should be interpreted as exploratory observations intended to guide future mechanistic investigations rather than establish definitive causal relationships.

### 2.2 Study Population

A total of twelve adult volunteers were enrolled in this exploratory study after providing written informed consent. The study population consisted of two groups. The HIV group included six untreated individuals with confirmed HIV-1 infection who had not previously received antiretroviral therapy. The control group comprised six HIV-seronegative healthy volunteers without evidence of chronic viral infection or known immunological disorders. Participants were recruited according to predefined eligibility criteria. Individuals with diabetes mellitus, chronic liver disease, chronic kidney disease, malignancy, autoimmune disorders, or other conditions known to influence immune function were excluded. Subjects receiving immunosuppressive medications or previous antiretroviral treatment were likewise excluded to minimize potential confounding effects on lymphocyte phenotype and function. Peripheral venous blood samples were collected under standardized conditions from all participants and processed immediately following collection. Identical laboratory procedures were applied to both study groups throughout the investigation to ensure analytical consistency.

### 2.3 Blood Collection and Sample Processing

Peripheral venous blood (10 mL) was collected from each participant by standard venipuncture into sterile ethylenediaminetetraacetic acid (EDTA)-anticoagulated collection tubes. Samples were processed immediately after collection to minimize ex vivo alterations in lymphocyte viability and phenotype. Whole blood aliquots were used directly for flow cytometric immunophenotyping and stimulation assays, whereas additional samples underwent density separation and cellular purification for downstream immunofluorescence imaging, ELISPOT analysis, and conjugation experiments. All laboratory procedures were performed under identical experimental conditions for both HIV-positive and control samples.

### 2.4 Flow Cytometric Immunophenotyping

Baseline immunophenotyping was performed using multiparameter flow cytometry to quantify circulating CD4□ and CD8□ T-lymphocyte populations in both study groups. Whole blood aliquots (50 µL) were incubated with fluorochrome-conjugated monoclonal antibodies directed against human CD4 and CD8 surface antigens according to the manufacturer’s recommendations. Following antibody incubation, erythrocytes were lysed, and leukocytes were suspended in phosphate-buffered saline before acquisition. Samples were analyzed using a calibrated flow cytometer under standardized acquisition settings. Absolute CD4□ and CD8□ T-cell counts were recorded for each participant and expressed as cells/µL of peripheral blood. These baseline measurements served as the reference values for all subsequent stimulation experiments.

### 2.5 Recombinant HIV-1 p17 Stimulation Assay

To investigate whether exposure to a structural HIV protein induces measurable alterations in T-cell phenotype, whole blood samples were incubated ex vivo with purified recombinant HIV-1 p17 matrix protein. Recombinant p17 was selected because of its documented immunomodulatory properties and reported interactions with multiple immune-cell populations. Aliquots of peripheral blood were incubated with recombinant p17 under standardized laboratory conditions and analyzed after 1, 2, 3, and 4 hours of exposure. Parallel control samples from healthy donors underwent identical incubation procedures. Following each incubation interval, samples were analyzed by flow cytometry to determine dynamic changes in CD4□ and CD8□ T-cell populations relative to baseline values. These experiments were designed to evaluate whether recombinant p17 exposure was associated with reproducible alterations in lymphocyte immunophenotype under controlled ex vivo conditions.

### 2.6 Isolation of CD4□ and CD8□ T-Cell Populations

To further investigate the immunophenotypic alterations observed following recombinant HIV-1 p17 stimulation, CD4□ and CD8□ T-cell populations were isolated from peripheral blood for subsequent immunofluorescence analysis. Cell purification was performed to enable direct visualization of phenotypic changes occurring at the individual-cell level while minimizing interference from other circulating leukocyte populations. Peripheral blood mononuclear cells were prepared from freshly collected blood samples following erythrocyte lysis using ammonium chloride lysis buffer. After repeated washing with phosphate-buffered saline (PBS) containing 1% bovine serum albumin (BSA), CD4□ and CD8□ T lymphocytes were isolated using magnetic bead–based immunoselection according to the manufacturer’s protocol. To reduce contamination by CD14-expressing monocytes, an initial monocyte depletion step was performed before T-cell purification. Purified lymphocyte populations were subsequently resuspended in autologous plasma and used immediately for downstream functional experiments.

### 2.7 Immunofluorescence Assessment of Cellular Phenotype

To investigate whether recombinant HIV-1 p17 stimulation was associated with alterations in T-cell immunophenotype, purified lymphocyte populations were examined using fluorescence immunolabeling. Purified CD4□ and CD8□ T-cell suspensions were incubated with recombinant HIV-1 p17 under standardized laboratory conditions. Fluorescent monoclonal antibodies directed against CD4 and CD8 surface antigens were subsequently added according to the experimental design. Samples were examined at predefined incubation intervals (1, 2, 3, and 4 h) using fluorescence microscopy. Particular attention was directed toward identifying cells exhibiting simultaneous immunoreactivity for both CD4- and CD8-associated surface markers. Representative microscopic fields were recorded for qualitative comparison between HIV-positive and control samples.

## 3. Results

### 3.1 Baseline Immunophenotypic Characteristics

To establish the immunological characteristics of the study population before experimental stimulation, baseline CD4□ and CD8□ T-lymphocyte counts were determined in peripheral blood obtained from untreated HIV-1-positive individuals and healthy controls using flow cytometric immunophenotyping. As expected, untreated HIV-positive participants demonstrated markedly lower circulating CD4□ T-cell counts than healthy controls, accompanied by a corresponding expansion of the CD8□ T-cell compartment. Despite these reciprocal changes, the combined CD4□/CD8□ T-cell count remained relatively stable between the two groups, suggesting that the overall T-lymphocyte population was preserved while its phenotypic distribution differed substantially. These baseline observations established the reference framework for all subsequent stimulation experiments and confirmed the characteristic immunophenotypic profile associated with chronic untreated HIV infection **Figure 1**.

**Figure 1.**
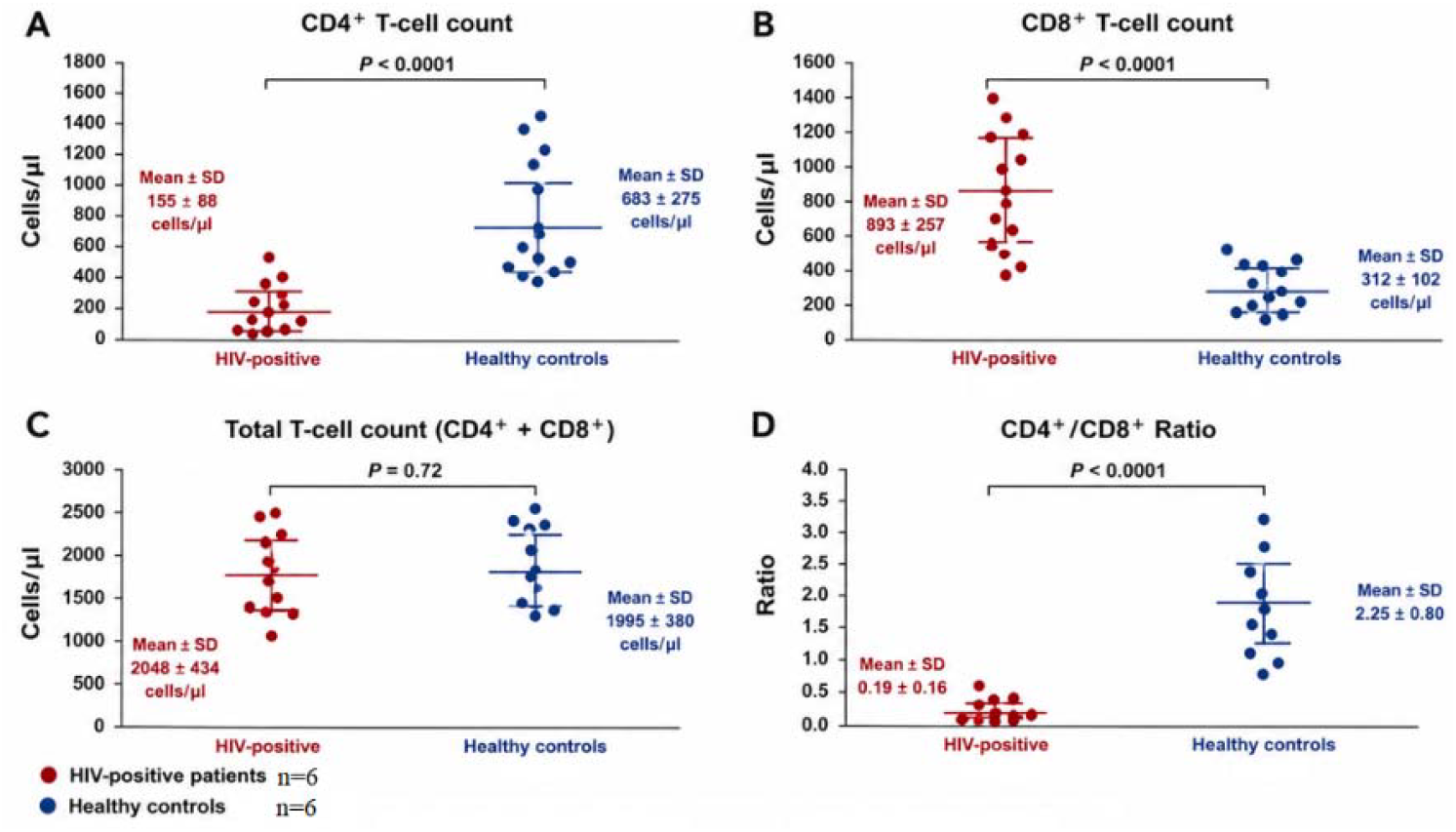
Baseline immunophenotypic characteristics of untreated HIV-positive participants and healthy controls. (A) Absolute CD4□ T-cell counts, (B) Absolute CD8□ T-cell counts, (C) Total circulating T-cell counts (CD4□+ CD8□) and (D) CD4□/CD8□ratio. Each point represents one individual participant. Horizontal bars indicate the mean ± SD. Statistical comparisons were performed using Welch’s t-test.

### 3.2 Dynamic Immunophenotypic Changes Following Recombinant HIV-1 p17 Exposure

To investigate whether exposure to recombinant HIV-1 p17 influenced T-cell phenotype, peripheral blood samples obtained from HIV-positive and healthy participants were incubated ex vivo with recombinant p17 and analyzed sequentially by flow cytometry after 1, 2, 3, and 4 hours of stimulation. In HIV-positive samples, recombinant p17 exposure was associated with a progressive reduction in the detectable CD4□ T-cell population together with a corresponding increase in CD8□ T-cell counts throughout the incubation period. These changes became increasingly apparent during prolonged incubation and reached their greatest magnitude after 4 hours of stimulation. In contrast, lymphocyte populations obtained from healthy controls remained remarkably stable throughout the same experimental period, demonstrating minimal variation in either CD4□or CD8□ T-cell counts.

Interestingly, although reciprocal alterations were observed in individual lymphocyte subsets, the total combined CD4□/CD8□ T-cell count remained relatively unchanged. This finding suggested that the observed changes reflected redistribution of immunophenotypic characteristics rather than generalized lymphocyte loss under the experimental conditions Figure 2,3.

**Figure 2.**
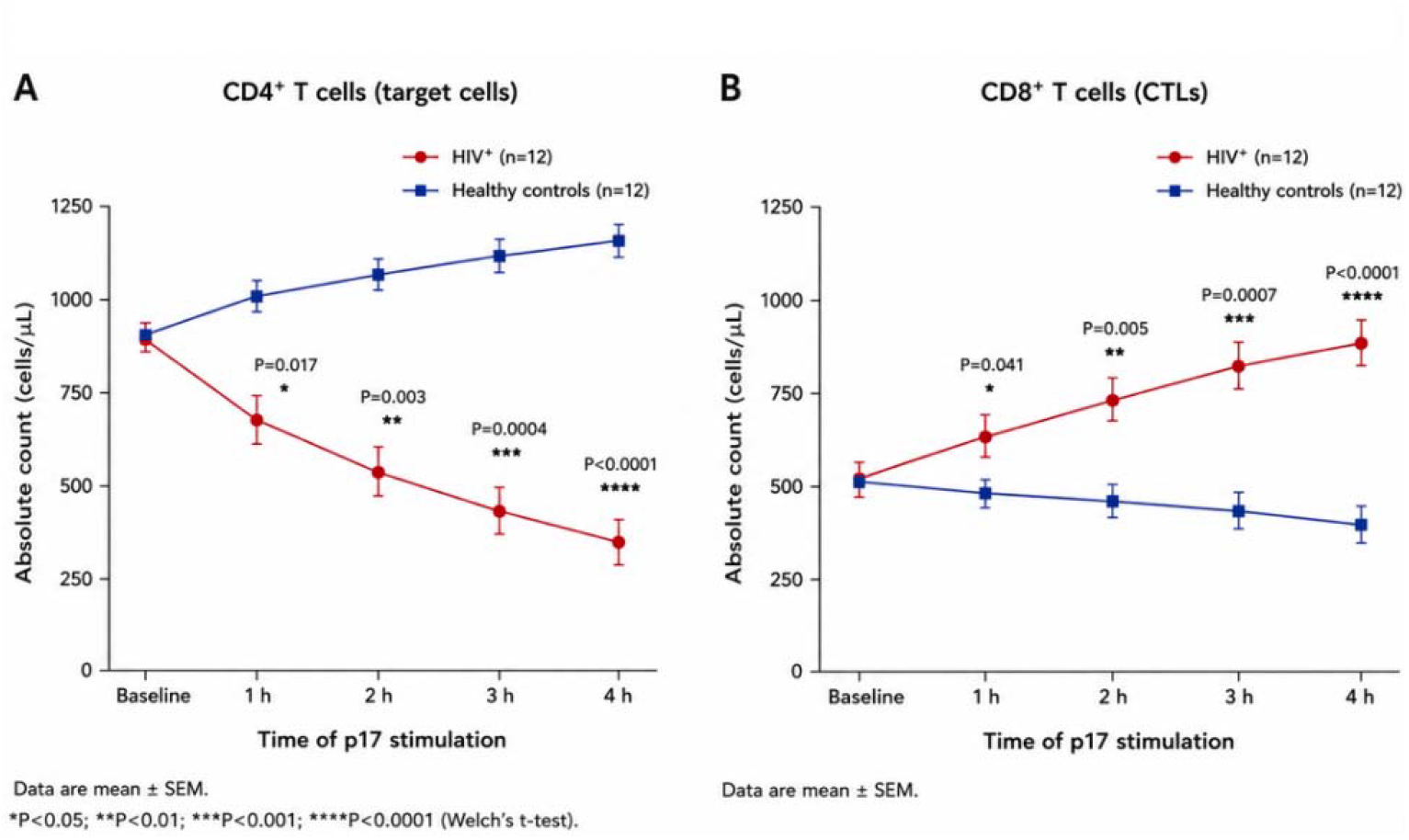
Time-dependent changes in CD4□ and CD8□ T-cell populations following ex vivo stimulation with recombinant HIV-1 p17 matrix protein. (A) Sequential changes in CD4□ T-cell counts during 1–4 h of recombinant HIV-1 p17 stimulation, (B) Sequential changes in CD8□ T-cell counts during the same incubation period. Thin lines represent individual participants, whereas bold lines and error bars represent group mean ± SD. Measurements were obtained by flow cytometric immunophenotyping.

**Figure 3.**
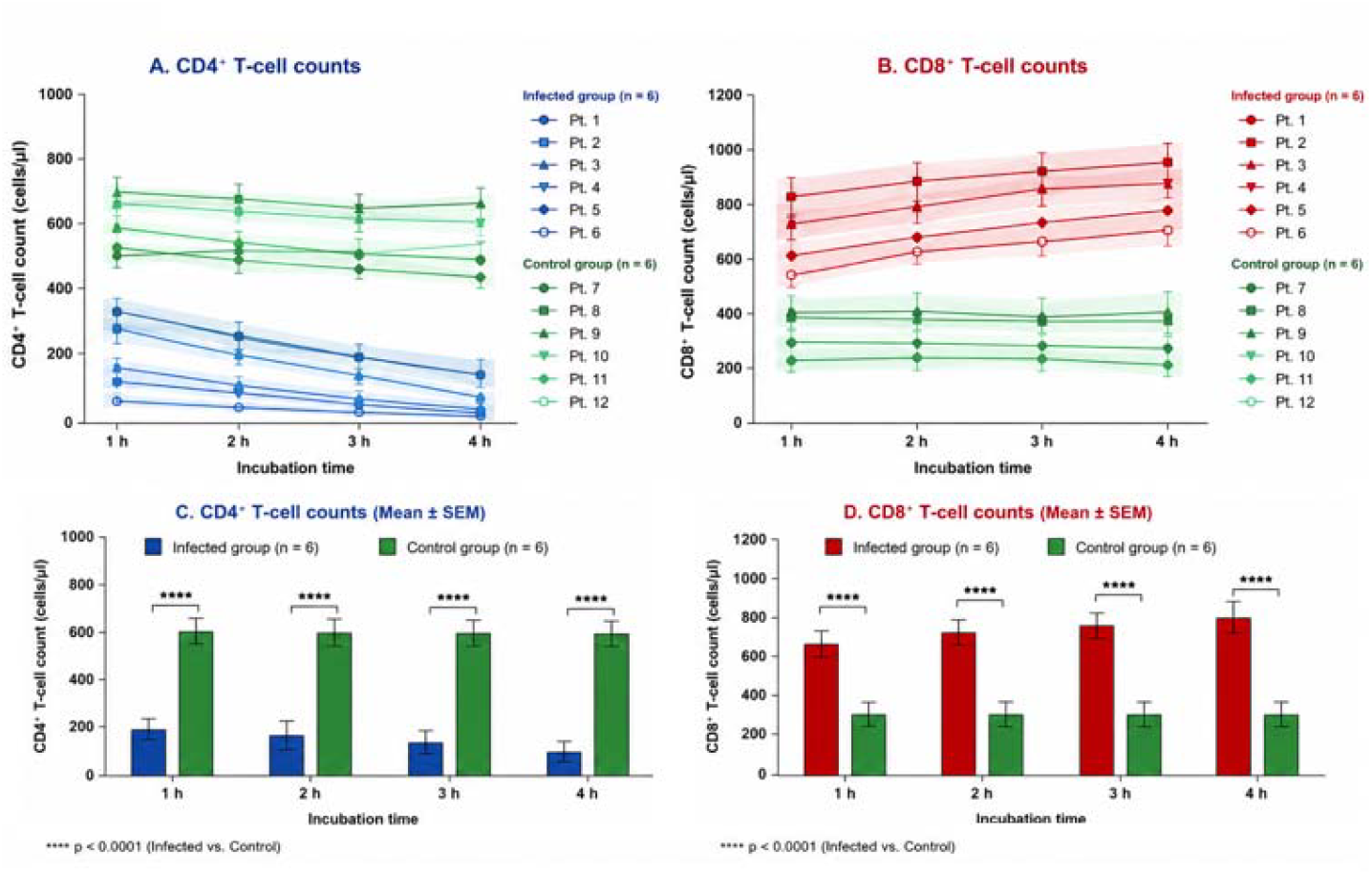
Relative preservation of the total circulating T-cell compartment during recombinant HIV-1 p17 stimulation. Changes in total circulating T-cell counts (CD4□+ CD8□) following ex vivo stimulation with recombinant HIV-1 p17 matrix protein. Despite reciprocal alterations in CD4□ and CD8□ T-cell populations, the overall T-cell count remained comparatively stable throughout the experimental period.

### 3.3 Immunofluorescence Evidence of Phenotypic Reprogramming

To further investigate the cellular basis of the immunophenotypic changes observed during recombinant HIV-1 p17 stimulation, purified lymphocyte populations were examined using dual-color immunofluorescence microscopy. Before stimulation, CD4□ and CD8□ T lymphocytes demonstrated the expected mutually exclusive immunolabeling patterns, with individual cells expressing either CD4- or CD8-associated surface markers. Following prolonged recombinant p17 exposure, however, a distinct cellular population exhibiting simultaneous immunoreactivity for both CD4 and CD8 markers became detectable in samples obtained from HIV-positive participants. The appearance of these dual-labeled cells was time dependent and became progressively more evident after extended incubation. Such cells were rarely observed in healthy control samples processed under identical experimental conditions. Representative fluorescence micrographs illustrating these observations are presented in **Figure 4**.

**Figure 4.**
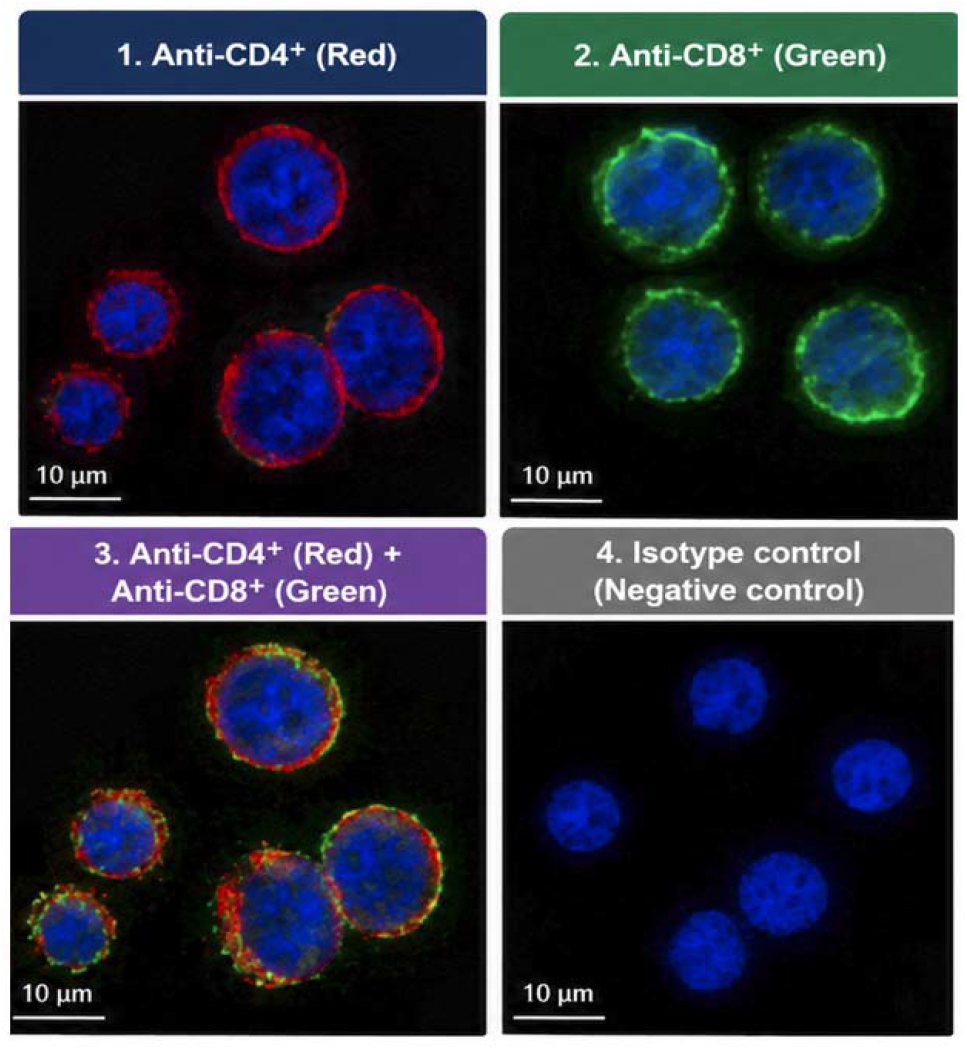
Immunofluorescence analysis of CD4□ and CD8□surface-marker localization following ex vivo stimulation with recombinant HIV-1 p17 matrix protein. (A) Representative fluorescence micrograph showing immunolabeling of CD4-associated surface markers (red) on CD4□ T cells. Cell nuclei were counterstained with DAPI (blue), (B) Representative fluorescence micrograph showing immunolabeling of CD8-associated surface markers (green) on CD8□ T cells. Cell nuclei were counterstained with DAPI (blue), (C) Merged fluorescence image demonstrating cells exhibiting simultaneous CD4-associated (red) and CD8-associated (green) immunoreactivity, resulting in areas of yellow/orange signal consistent with co-localization of both markers following recombinant HIV-1 p17 stimulation, (D) Representative negative (isotype) control showing the absence of specific immunofluorescent staining, confirming the specificity of the antibody labeling under the experimental conditions. **Scale bars = 10** μ**m. Images are representative of repeated experiments performed under identical laboratory conditions**.

### 3.4 Immunophenotypic Evidence of Cellular Reprogramming

Purified lymphocyte populations obtained from HIV-positive participants were subsequently examined by dual-color immunofluorescence microscopy to investigate whether the reciprocal alterations observed by flow cytometry were accompanied by changes in surface-marker expression at the individual-cell level. Before recombinant HIV-1 p17 stimulation, CD4□ and CD8□lymphocytes demonstrated the expected mutually exclusive staining pattern, with individual cells expressing either CD4-associated or CD8-associated surface markers. Following prolonged exposure to recombinant p17, however, an additional cellular population exhibiting simultaneous immunoreactivity for both CD4 and CD8 markers became detectable. The appearance of these dual-labeled cells occurred progressively during incubation and was consistently more apparent in samples obtained from HIV-positive individuals than in healthy controls processed under identical experimental conditions. Representative immunofluorescence micrographs illustrating these observations are presented in **Figure 5**.

**Figure 5.**
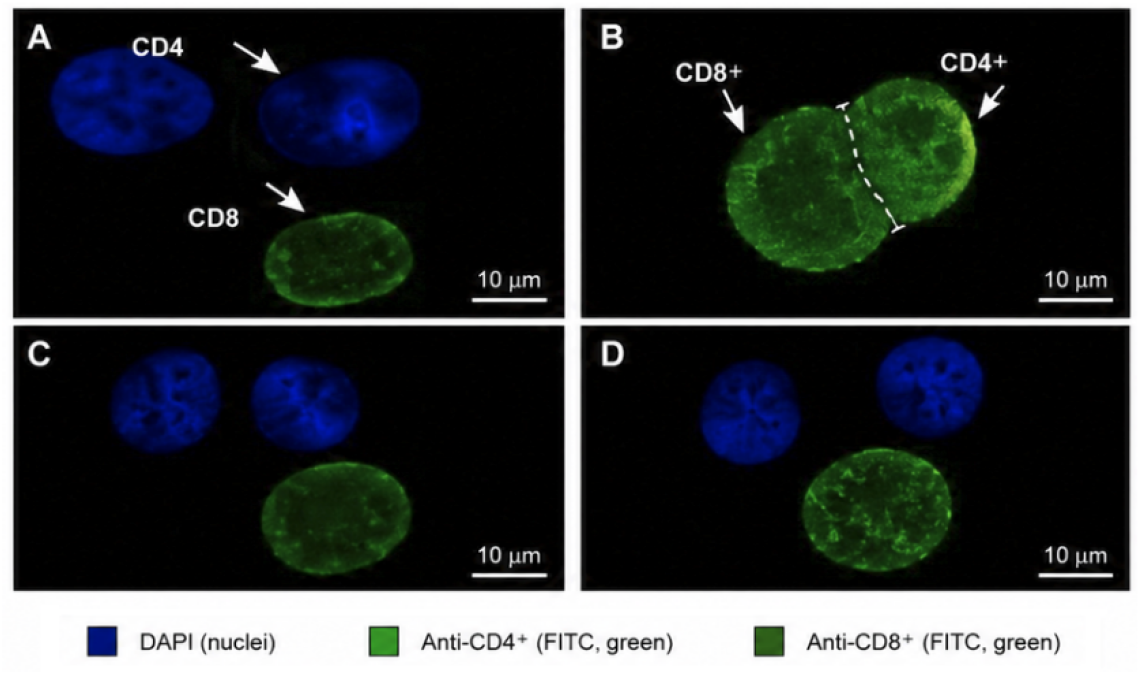
Conjugate formation assay demonstrating cellular interaction patterns between CD8□ T cells and CD4□ target cells following ex vivo recombinant HIV-1 p17 stimulation. **(A)** Representative fluorescence micrograph showing a mixed-cell preparation before stable cell-cell contact. CD4□ Target cells (blue fluorescence) and CD8□ T cells (green fluorescence) are visualized before conjugate formation, **(B)** Representative image demonstrating conjugate formation between a CD8□ T cell and a CD4□ Target cell following recombinant HIV-1 p17 stimulation. The close membrane-to-membrane contact illustrates stable cellular interaction under the experimental conditions, **(C)** Representative suspended-cell preparation in which CD4□ Target cells and CD8□ T cells remain spatially separated, with no evidence of stable conjugate formation, **(D)** Representative suspended-cell mixture following recombinant HIV-1 p17 stimulation showing the absence of stable cellular contact. Under these conditions, CD8□ T cells did not exhibit acquisition of detectable CD4-associated surface immunolabeling, Images are representative of repeated experiments performed under identical laboratory conditions. The conjugation assay was designed to evaluate changes in cellular interaction patterns and does not by itself establish the molecular mechanism responsible for the observed interactions.

The detection of cells expressing both CD4- and CD8-associated markers does not by itself establish lineage conversion or irreversible cellular transformation. However, these observations indicate the emergence of an altered immunophenotypic state following recombinant HIV-1 p17 stimulation and provided the rationale for subsequent functional characterization of this cellular population.

### 3.5 Functional Characterization of Cells Exhibiting Altered Immunophenotypic Characteristics

To determine whether the altered immunophenotypic profile observed following recombinant HIV-1 p17 stimulation was accompanied by functional changes, cytokine-producing capacity was evaluated using an interferon-γ (IFN-γ) enzyme-linked immunospot (ELISPOT) assay. Purified lymphocyte populations displaying altered immunophenotypic characteristics following recombinant p17 exposure demonstrated a marked reduction in IFN-γ secretion when compared with conventionally identified CD8□ T cells obtained from healthy control samples. In contrast, CD8□ T cells isolated from healthy individuals exhibited robust cytokine production comparable to the positive assay controls, confirming preserved functional activity under identical experimental conditions. The reduction in IFN-γ production observed in the altered cell population suggests that acquisition of modified surface-marker expression was accompanied by measurable impairment of cellular effector function. Although the present experiments were not designed to identify the molecular mechanisms responsible for this functional alteration, the findings indicate that the observed immunophenotypic changes were associated with biological consequences extending beyond simple alterations in receptor expression. Collectively, these observations support the concept that recombinant HIV-1 p17 stimulation was associated with the emergence of a lymphocyte population displaying both altered immunophenotypic characteristics and reduced functional responsiveness under the experimental conditions employed **Figure 6**.

**Figure 6.**
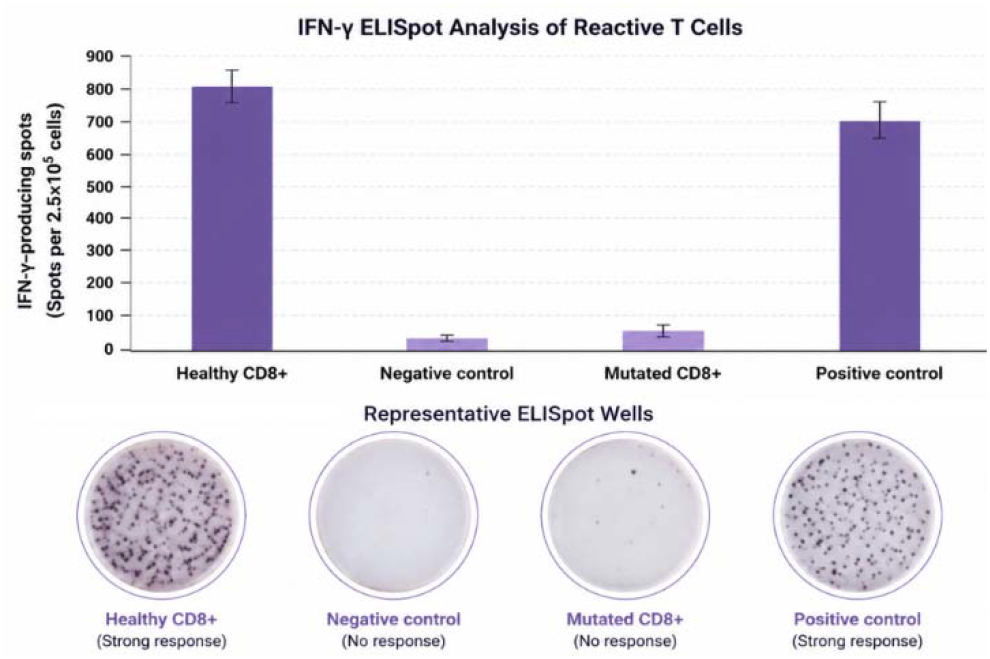
Immunofluorescence analysis of CD4□ and CD8□surface-marker localization following ex vivo stimulation with recombinant HIV-1 p17 matrix protein. (A) Representative fluorescence micrograph showing immunolabeling of CD4-associated surface markers (red) on CD4□ T cells. Cell nuclei were counterstained with DAPI (blue), (B) Representative fluorescence micrograph showing immunolabeling of CD8-associated surface markers (green) on CD8□ T cells. Cell nuclei were counterstained with DAPI (blue), (C) Merged fluorescence image demonstrating cells exhibiting simultaneous CD4-associated (red) and CD8-associated (green) immunoreactivity, resulting in areas of yellow/orange signal consistent with co-localization of both markers following recombinant HIV-1 p17 stimulation, (D) Representative negative (isotype) control showing the absence of specific immunofluorescent staining, confirming the specificity of the antibody labeling under the experimental conditions.

### 3.6 Altered Cellular Interaction Patterns Following Recombinant HIV-1 p17 Exposure

To further investigate the biological behavior of cells exhibiting altered immunophenotypic characteristics, cellular interaction studies were performed using fluorescence-based conjugation assays. Purified CD8□ T cells isolated from HIV-positive participants formed conjugates with CD4□ T cells obtained from healthy donors under the experimental conditions employed. In contrast, comparable conjugate formation was not observed when CD8□ T cells from healthy controls were incubated with CD4□ T cells isolated from HIV-positive participants. These observations indicate that recombinant HIV-1 p17–associated phenotypic alterations were accompanied by measurable differences in cellular interaction patterns. Although the precise molecular basis of these altered interactions remains to be determined, the findings suggest that changes in immunophenotype may influence cell-to-cell communication in addition to modifying cytokine production. The present experiments were not designed to establish whether these altered interaction patterns directly contribute to HIV transmission or disease progression. Rather, they demonstrate that lymphocyte populations exhibiting altered immunophenotypic characteristics also display distinct functional behavior during controlled ex vivo cellular interaction assays.

### 3.7 Integrated Experimental Findings

Taken together, the experimental findings demonstrated a consistent sequence of immunological observations following ex vivo recombinant HIV-1 p17 stimulation. HIV-positive samples initially exhibited the expected reduction in circulating CD4□ T cells accompanied by expansion of the CD8□ T-cell compartment. Subsequent stimulation experiments revealed progressive alterations in these immunophenotypic profiles while preserving the overall size of the measured T-lymphocyte population. Immunofluorescence microscopy further identified cells displaying simultaneous CD4- and CD8-associated marker expression, suggesting the emergence of an altered immunophenotypic state following prolonged recombinant p17 exposure. Functional analyses demonstrated reduced IFN-γ production within this cellular population, whereas conjugation assays identified modified cellular interaction patterns under identical experimental conditions.

Individually, none of these observations establishes the mechanism responsible for the observed immunophenotypic alterations. Collectively, however, they support a coherent experimental model in which recombinant HIV-1 p17 exposure is associated with coordinated changes in lymphocyte phenotype and function.

## Discussion

The pathogenesis of human immunodeficiency virus type 1 (HIV-1) has been investigated extensively for more than four decades, leading to the establishment of several complementary mechanisms responsible for progressive immune dysfunction. Current models attribute the decline in circulating CD4□ T lymphocytes to a combination of direct viral cytopathic effects, chronic immune activation, activation-induced apoptosis, pyroptosis of abortively infected cells, immune-mediated clearance, and persistent inflammatory signaling **[5–11]**. Together, these mechanisms have substantially advanced our understanding of HIV-associated immunodeficiency and continue to form the foundation of current therapeutic strategies. Nevertheless, despite remarkable progress in defining these processes, important questions remain regarding the relationship between quantitative depletion of helper T cells and the profound functional impairment of cellular immunity that characterizes chronic HIV infection. Clinical observations demonstrating persistent immune dysregulation despite effective suppression of viral replication further suggest that additional biological mechanisms may contribute to disease progression **[2–4**,**19–23]**.

The present study explored this possibility by examining whether exposure to recombinant HIV-1 p17 matrix protein was associated with coordinated alterations in T-cell phenotype and function under controlled ex vivo conditions. Rather than focusing exclusively on numerical changes in lymphocyte populations, the study investigated whether prolonged viral protein stimulation produced qualitative alterations that could provide additional insight into HIV-associated immune dysfunction. Several consistent observations emerged from these experiments. Recombinant p17 stimulation produced progressive reductions in detectable CD4□ T-cell counts accompanied by reciprocal increases in CD8□ T-cell populations while preserving the overall size of the measured T-lymphocyte compartment. These quantitative findings were complemented by immunofluorescence observations demonstrating the appearance of cells exhibiting simultaneous CD4- and CD8-associated immunoreactivity, together with functional evidence showing reduced interferon-γ production and altered cellular interaction patterns. Although each observation individually provides only limited mechanistic information, their convergence suggests that recombinant HIV-1 p17 exposure is associated with coordinated alterations extending beyond simple quantitative changes in lymphocyte numbers **[32–38]**.

One possible biological interpretation of these findings is provided by the modern concept of immune-cell phenotypic reprogramming. During the past decade, increasing evidence has demonstrated that differentiated immune cells retain considerable functional plasticity and may undergo adaptive remodeling in response to prolonged inflammatory stimulation, chronic antigen exposure, or persistent cytokine signaling **[24–31]**. These adaptive processes involve coordinated alterations in transcriptional regulation, epigenetic programming, chromatin accessibility, metabolic activity, receptor expression, and cellular function without necessarily implying irreversible genetic mutation or complete lineage conversion **[24–31]**. Within this biological framework, the emergence of cells displaying altered immunophenotypic characteristics following recombinant HIV-1 p17 stimulation may reflect phenotypic remodeling induced by persistent viral signaling rather than permanent transformation of one lymphocyte lineage into another. Such an interpretation is consistent with contemporary understanding of immune-cell plasticity and provides a biologically plausible explanation for the experimental observations reported in the present study.

Recent evidence further indicates that not all CD4□ T-cell subsets contribute equally to HIV pathogenesis. Among these, T helper 17 (Th17) cells exhibit preferential susceptibility to HIV infection and play an essential role in maintaining gastrointestinal mucosal integrity and limiting microbial translocation. Early depletion and functional impairment of Th17 cells contribute to disruption of epithelial barrier function, persistent microbial translocation, systemic immune activation, and chronic inflammation, all of which are now recognized as central drivers of HIV disease progression **[45–47]**. Importantly, these observations demonstrate that qualitative functional alterations within specific CD4□ T-cell populations can profoundly influence immune homeostasis independently of absolute CD4□ T-cell numbers. Although the present study did not specifically investigate Th17 biology, the recognized functional plasticity and selective vulnerability of this subset provide additional biological plausibility for investigating immunophenotypic reprogramming as a complementary mechanism contributing to HIV-associated immune dysregulation.

The functional findings further strengthen this interpretation. The observed reduction in interferon-γ production indicates that the altered cell population was characterized not only by changes in surface-marker expression but also by measurable impairment of effector function. Similarly, the altered conjugation behavior observed during cellular interaction assays suggests that modifications in cellular phenotype may influence immune communication in addition to cytokine production. Together, these findings support the concept that chronic viral stimulation may induce coordinated functional remodeling affecting multiple aspects of lymphocyte biology simultaneously. However, the present experiments were not designed to determine the molecular pathways responsible for these changes, nor can they distinguish definitively between altered receptor expression, transcriptional reprogramming, selective expansion of previously existing subpopulations, or other mechanisms capable of producing similar experimental observations **[19–31**,**41–44]**.

Accordingly, the biological model proposed in this study should be regarded as hypothesis-generating rather than definitive. We propose that phenotypic reprogramming of a subset of CD4□ T lymphocytes may represent an additional mechanism contributing to HIV-associated immune dysregulation alongside the established processes of viral cytopathicity, apoptosis, pyroptosis, chronic immune activation, and immune exhaustion **[5–18]**. This model does not seek to replace current concepts of HIV immunopathogenesis but instead expands them by suggesting that qualitative alterations in lymphocyte phenotype and function may coexist with quantitative cellular depletion. Future studies employing lineage-tracing approaches, high-dimensional flow cytometry, single-cell transcriptomics, epigenomic profiling, proteomics, and metabolic analyses will be required to determine whether the altered immunophenotypic characteristics observed in the present study represent stable biological reprogramming, transient activation-dependent changes, or alternative mechanisms not evaluated within the current experimental design **[19–31]**.

Despite its exploratory nature, the present investigation possesses several strengths. Multiple complementary experimental approaches were used to evaluate both immunophenotypic and functional characteristics of lymphocyte populations, and the convergence of findings across independent methodologies increases confidence that the observed alterations are biologically meaningful. At the same time, several limitations should be acknowledged. The study involved a relatively small number of participants, employed ex vivo stimulation with a single recombinant HIV protein rather than complete viral infection, and did not incorporate molecular techniques capable of defining lineage relationships or transcriptional mechanisms. Consequently, the conclusions should be interpreted within the context of these methodological limitations.

## Conclusion

The present study provides experimental evidence that ex vivo stimulation with recombinant HIV-1 p17 matrix protein is associated with coordinated alterations in CD4□ and CD8□ T-cell immunophenotypes and function. The observed reciprocal changes in CD4□ and CD8□ T-cell populations, preservation of the overall T-cell compartment, emergence of cells exhibiting simultaneous CD4- and CD8-associated immunoreactivity, reduced interferon-γ production, and altered cellular interaction patterns collectively support the hypothesis that HIV-associated stimulation may promote **immunophenotypic reprogramming** of a subset of CD4□ T cells. Rather than challenging established mechanisms of HIV immunopathogenesis, the proposed model should be regarded as a complementary, hypothesis-generating framework that may help explain aspects of immune dysfunction not fully accounted for by quantitative CD4□ T-cell depletion alone. These findings provide a basis for further investigation into the contribution of qualitative immune-cell remodeling to chronic HIV-associated immune dysregulation.

## Future Directions

The observations reported in this study require independent validation using contemporary molecular and cellular technologies. Future investigations should combine high-dimensional flow cytometry, fluorescence-activated cell sorting, lineage-associated transcription factor analysis (including ThPOK **and** Runx3), single-cell RNA sequencing, T-cell receptor clonotyping, epigenomic profiling, and proteomic analyses to determine whether the altered immunophenotypic characteristics identified in the present work represent stable cellular reprogramming, transient activation-dependent remodeling, or alternative biological processes.

Additional studies employing replication-competent HIV infection models, larger patient cohorts, and longitudinal clinical follow-up will also be essential to determine the biological significance and potential clinical relevance of immunophenotypic reprogramming in HIV-associated immune dysfunction. A clearer understanding of these mechanisms may ultimately contribute to the identification of novel biomarkers and immunomodulatory therapeutic strategies aimed at restoring immune homeostasis in people living with HIV.

## Ethics Statement

Peripheral blood samples were obtained from adult participants after written informed consent had been obtained prior to sample collection. All samples were anonymized before laboratory analysis to ensure participant confidentiality. The study was conducted in accordance with the ethical principles of the Declaration of Helsinki governing research involving human participants. The research was performed using samples collected at a private hospital, where formal institutional ethics committee review was not available at the time of sample collection.

## Author Contributions

**Conceptualization:** Sherif Salah and Khalid F. Kassem; **Methodology:** Sherif Salah and Mohamed Sherif; **Investigation:** Sherif Salah; **Data Curation:** Sherif Salah and Mohamed Sherif; **Formal Analysis:** Sherif Salah and Khalid F. Kassem; **Visualization:** Sherif Salah; **Writing Original Draft Preparation:** Sherif Salah; **Writing—Review and Editing:** Khalid F. Kassem, Mohamed Sherif, and Ashraf M. Abu-Seida; **Supervision:** Khalid F. Kassem and Ashraf M. Abu-Seida; **Project Administration:** Sherif Salah. All authors have read and agreed to the published version of the manuscript.

## Data Availability Statement

The Data Availability Statement has been revised to provide complete information regarding the public availability of the datasets supporting this study. Raw datasets, Supplementary Figures, are publicly available through the Zenodo repository: https://doi.org/10.5281/zenodo.21428540

