## Supplementary S1 for "Experimental Evidence for HIV-Associated Phenotypic Reprogramming of CD4□ T Cells: An Exploratory Immunological Study"

Supplementary S1 (Cover Figure): Conceptual illustration of the proposed mechanism investigated in this study. Ex vivo stimulation with recombinant HIV-1 p17 was associated with coordinated immunophenotypic and functional alterations in CD4⁺ T cells, including altered CD4/CD8-associated marker localization, impaired cellular interactions, and reduced immune function. These findings support a hypothesis of HIV-associated immunophenotypic reprogramming as a potential contributor to chronic immune dysregulation.

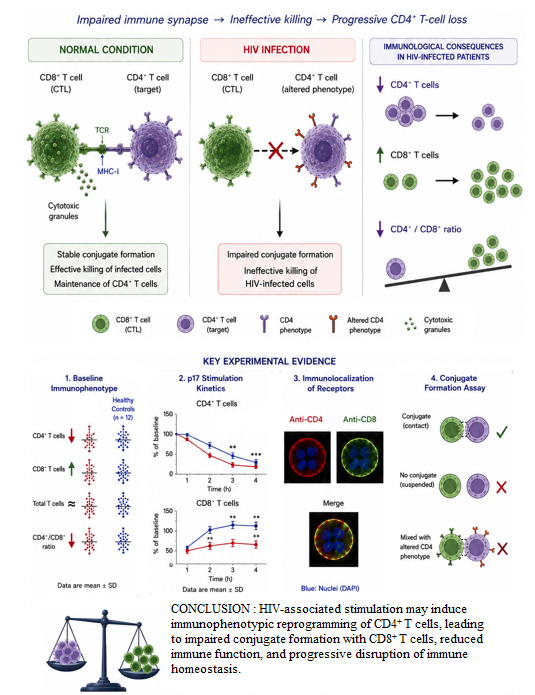

**Figure 1. Baseline immunophenotypic characteristics of untreated HIV-positive participants and healthy controls.** (A) Absolute CD4⁺ T-cell counts, (B) Absolute CD8⁺ T-cell counts, (C) Total circulating T-cell counts (CD4⁺ + CD8⁺) and (D) CD4⁺/CD8⁺ ratio. Each point represents one individual participant. Horizontal bars indicate the mean ± SD. Statistical comparisons were performed using Welch's t-test.

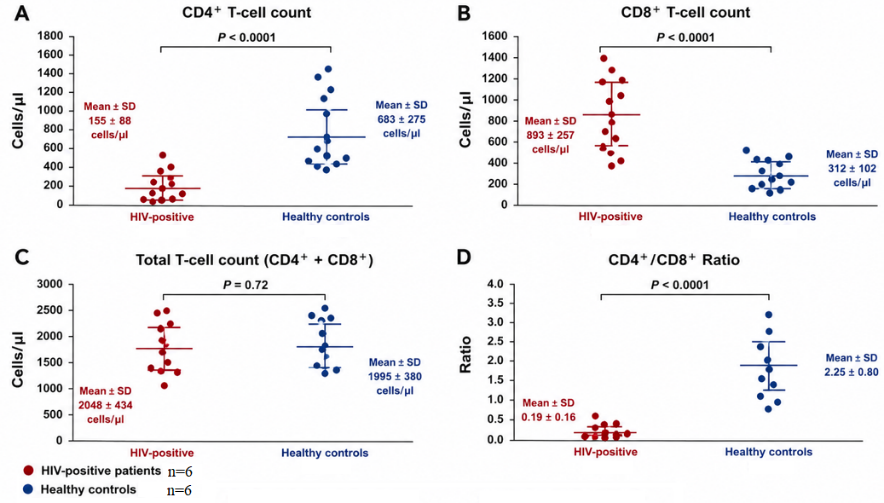

**Figure 2. Time-dependent changes in CD4⁺ and CD8⁺ T-cell populations following ex vivo stimulation with recombinant HIV-1 p17 matrix protein.** (A) Sequential changes in CD4⁺ T-cell counts during 1–4 h of recombinant HIV-1 p17 stimulation, (B) Sequential changes in CD8⁺ T-cell counts during the same incubation period. Thin lines represent individual participants, whereas bold lines and error bars represent group mean ± SD. Measurements were obtained by flow cytometric immunophenotyping.

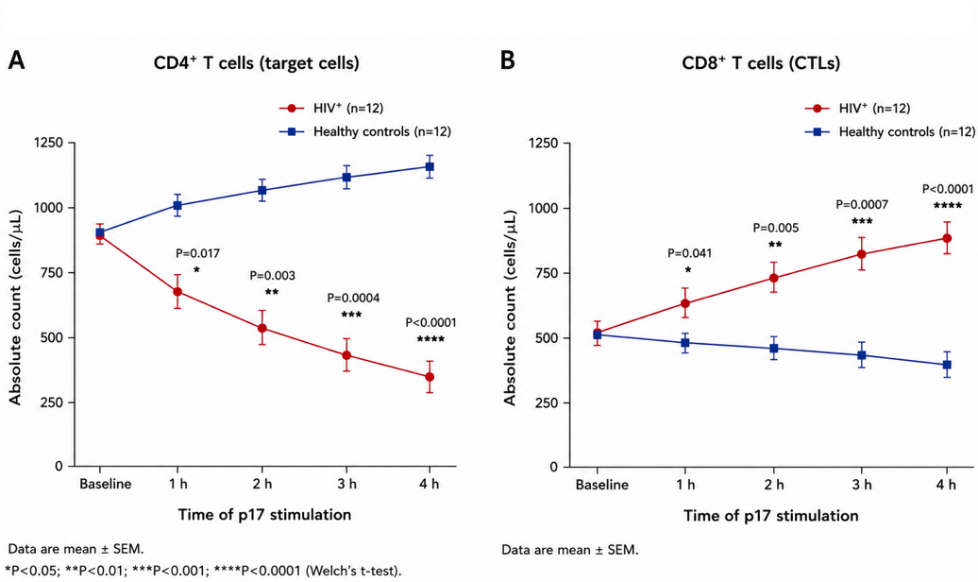

**Figure 3. Relative preservation of the total circulating T-cell compartment during recombinant HIV-1 p17 stimulation.** Changes in total circulating T-cell counts (CD4⁺ + CD8⁺) following ex vivo stimulation with recombinant HIV-1 p17 matrix protein. Despite reciprocal alterations in CD4⁺ and CD8⁺ T-cell populations, the overall T-cell count remained comparatively stable throughout the experimental period.

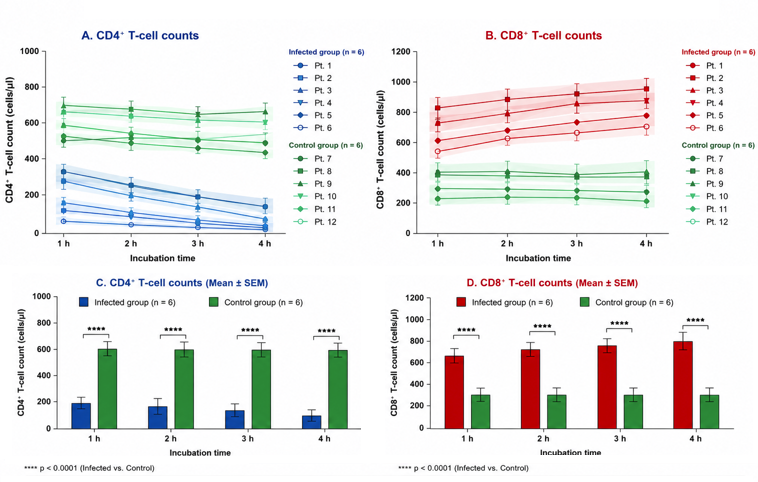

**Figure 4. Functional assessment of lymphocyte activity by interferon-γ ELISPOT analysis.** Interferon-γ-producing cells were quantified following ex vivo stimulation with recombinant HIV-1 p17 matrix protein. Healthy CD8⁺ T cells demonstrated robust interferon-γ secretion comparable to the positive control, whereas cells exhibiting altered immunophenotypic characteristics showed markedly reduced cytokine production, like the negative control. Representative ELISPOT wells are shown below the quantitative analysis. **Scale bars = 10 μm. Images are representative of repeated experiments performed under identical laboratory conditions.**

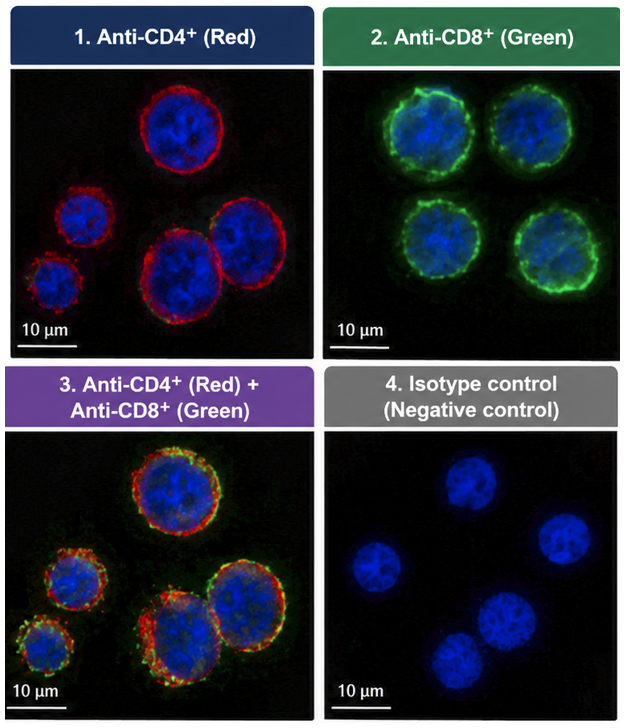

**Figure 5. Conjugate formation assay demonstrating cellular interaction patterns between CD8⁺ T cells and CD4⁺ target cells following ex vivo recombinant HIV-1 p17 stimulation. (A)** Representative fluorescence micrograph showing a mixed-cell preparation before stable cell-cell contact. CD4⁺ target cells (blue fluorescence) and CD8⁺ T cells (green fluorescence) are visualized before conjugate formation, **(B)** Representative image demonstrating conjugate formation between a CD8⁺ T cell and a CD4⁺ target cell following recombinant HIV-1 p17 stimulation. The close membrane-to-membrane contact illustrates stable cellular interaction under the experimental conditions, **(C)** Representative suspended-cell preparation in which CD4⁺ target cells and CD8⁺ T cells remain spatially separated, with no evidence of stable conjugate formation, **(D)** Representative suspended-cell mixture following recombinant HIV-1 p17 stimulation showing the absence of stable cellular contact. Under these conditions, CD8⁺ T cells did not exhibit acquisition of detectable CD4-associated surface immunolabeling,

Images are representative of repeated experiments performed under identical laboratory conditions. The conjugation assay was designed to evaluate changes in cellular interaction patterns and does not by itself establish the molecular mechanism responsible for the observed interactions.

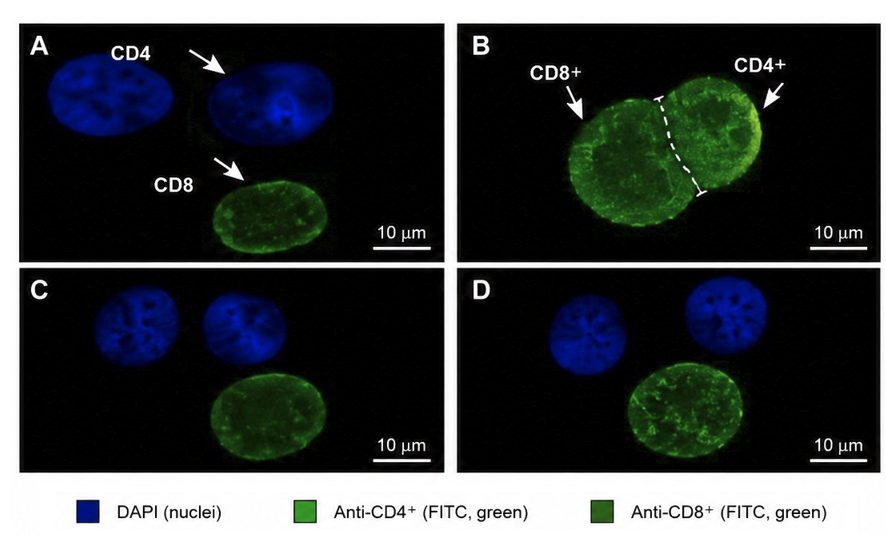

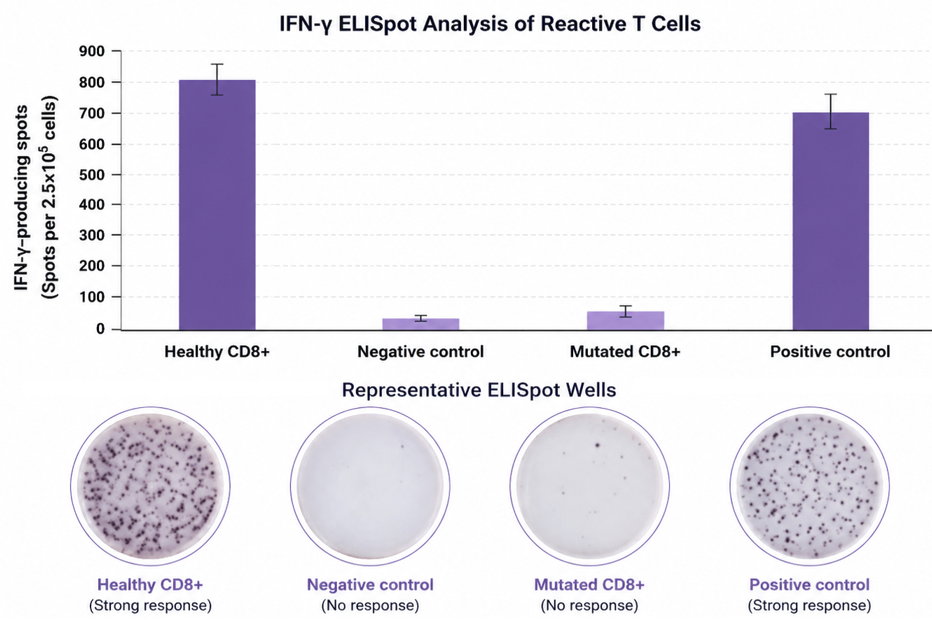

### ****Figure 6. Immunofluorescence analysis of CD4⁺ and CD8⁺ surface-marker localization following ex vivo stimulation with recombinant HIV-1 p17 matrix protein. (A)**** Representative fluorescence micrograph showing immunolabeling of CD4-associated surface markers (red) on CD4⁺ T cells. Cell nuclei were counterstained with DAPI (blue), ****(B)**** Representative fluorescence micrograph showing immunolabeling of CD8-associated surface markers (green) on CD8⁺ T cells. Cell nuclei were counterstained with DAPI (blue), ****(C)**** Merged fluorescence image demonstrating cells exhibiting simultaneous CD4-associated (red) and CD8-associated (green) immunoreactivity, resulting in areas of yellow/orange signal consistent with co-localization of both markers following recombinant HIV-1 p17 stimulation, ****(D)**** Representative negative (isotype) control showing the absence of specific immunofluorescent staining, confirming the specificity of the antibody labeling under the experimental conditions.
